# Investigation of COVID-19 comorbidities reveals genes and pathways coincident with the SARS-CoV-2 viral disease

**DOI:** 10.1101/2020.09.21.306720

**Authors:** Mary E. Dolan, David P. Hill, Gaurab Mukherjee, Monica S. McAndrews, Elissa J. Chesler, Judith A. Blake

## Abstract

The emergence of the SARS-CoV-2 virus and subsequent COVID-19 pandemic initiated intense research into the mechanisms of action for this virus. It was quickly noted that COVID-19 presents more seriously in conjunction with other human disease conditions such as hypertension, diabetes, and lung diseases. We conducted a bioinformatics analysis of COVID-19 comorbidity-associated gene sets, identifying genes and pathways shared among the comorbidities, and evaluated current knowledge about these genes and pathways as related to current information about SARS-CoV-2 infection. We performed our analysis using GeneWeaver (GW), Reactome, and several biomedical ontologies to represent and compare common COVID-19 comorbidities. Phenotypic analysis of shared genes revealed significant enrichment for immune system phenotypes and for cardiovascular-related phenotypes, which might point to alleles and phenotypes in mouse models that could be evaluated for clues to COVID-19 severity. Through pathway analysis, we identified enriched pathways shared by comorbidity datasets and datasets associated with SARS-CoV-2 infection.

## Introduction

COVID-19 emerged as a global pandemic through the first half of 2020^1^. The severity of the disease varies from asymptomatic to lethal with a case mortality rate in the 20 most affected countries ranging between 1 and 15.3% (https://coronavirus.jhu.edu/data/mortality ; retrieved 24 July 2020). Severe disease shows manifestations of both acute respiratory distress syndrome (ARDS) and cytokine release syndrome (CRS)^2,3^. In pediatric patients, a blood vessel inflammatory pathology similar to Kawasaki disease is sometimes present^4^. All of these presentations have common elements of abnormality of inflammatory responses and manifestations of vascular defects such as thrombosis, which may be causally related^5,6,7,8,9^.

Since the emergence and global transmission of the SARS-CoV-2 virus, many studies have reported that patients with certain underlying medical conditions have especially severe responses to the coronavirus infection^10^. Some of the identified comorbidities that lead to severe disease are Cardiovascular Disease, Diabetes, Hepatitis, Lung Disease, and Kidney Disease^11,12,13,14,15,16,17^.

Understanding what makes some patients suffer from severe COVID-19 is an ongoing puzzle that is being investigated from both the virus and host perspectives^18^. We hypothesize that by exploring the underlying genetic basis of comorbidities associated with severe disease, we can identify putative host genes and pathways that are responsible for or contribute to the severity. Identification of these genes and pathways can serve as a gateway for further investigation into understanding how the host responds to the virus and for potential therapeutic strategies to interfere with a severe outcome.

We interrogated gene sets that are associated with the five previously mentioned underlying comorbidities to determine gene products that are shared among them. We identified several pathways and phenotypes in common, including those that are associated with severe COVID-19 pathology. All of the comorbid diseases have been and continue to be actively studied, now in the additional context of response to SARS-CoV-2 infection^19,20,21,22,23^. In particular, the laboratory mouse has been extensively utilized as an animal model to study these conditions^24^. As a result, mouse strains carrying mutations in shared genes or genes in shared pathways, and engineered to be capable of being infected by the virus, can present useful starting points for investigating the biological basis of disease severity^25^.

We report here on investigations of the host genetics and genomics of a set of comorbidity conditions. We include data identifying the shared pathways and cellular mechanisms associated with these diseases and correlate these data with recent studies of the genetic basis of the COVID-19 to identify elements that are shared among comorbidities and the host response to the disease. Our results suggest specific directions of future study to understand the genetic foundation of severe COVID-19.

## Materials and Methods

### Gene Sets Used for Analysis

All gene sets used in our analysis are publicly available from the GeneWeaver resource (www.geneweaver.org)^26^. Genes associated with Cardiovascular Disease, Diabetes, Hepatitis, Lung Disease were derived from gene sets associated with MeSH terms that relate to these comorbidities. The gene set for Kidney Disease was derived from the union of genes associated with Proteinuria Hematuria, Elevated Serum Creatinine, Increased Blood Urea Nitrogen and Decreased Glomerular Filtration Rate in the Human Phenotype Ontology (HPO)^27^. The genes in the MeSH and HPO gene sets and associated metadata (indicating their association with COVID-19 and citations supporting the association) were incorporated into GW and used for analysis. The comorbidity-related gene sets are shown in Table 1.

**Table 1:**
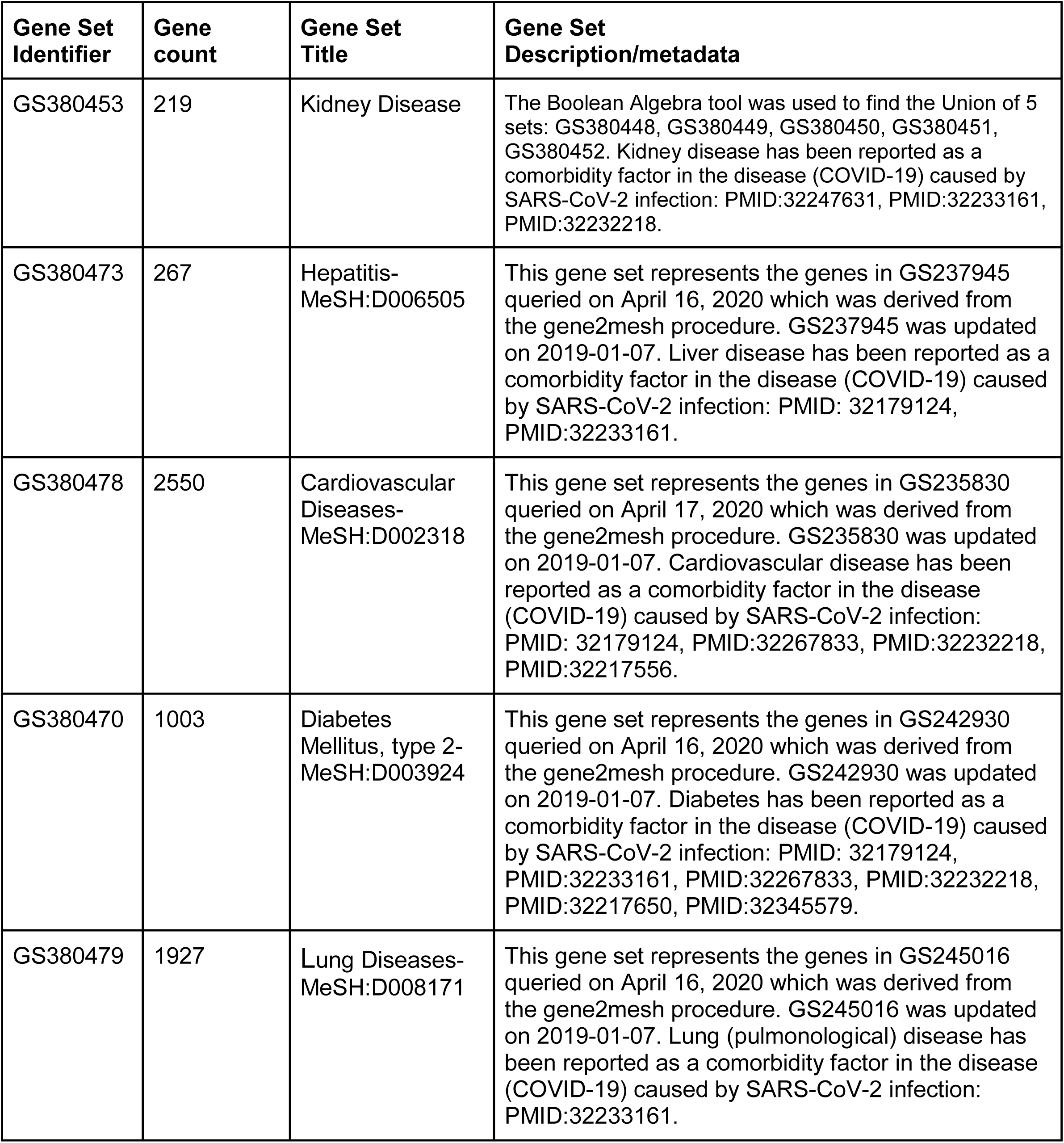
SARS-CoV-2 (COVID-19) comorbidity-related gene sets

| **Gene Set Identifier** | **Gene count** | **Gene Set Title** | **Gene Set Description/metadata** |
| --- | --- | --- | --- |
| GS380453 | 219 | Kidney Disease | The Boolean Algebra tool was used to find the Union of 5 sets: GS380448, GS380449, GS380450, GS380451, GS380452. Kidney disease has been reported as a comorbidity factor in the disease (COVID-19) caused by SARS-CoV-2 infection: PMID:32247631, PMID:32233161, PMID:32232218. |
| GS380473 | 267 | Hepatitis-MeSH:D006505 | This gene set represents the genes in GS237945 queried on April 16, 2020 which was derived from the gene2mesh procedure. GS237945 was updated on 2019-01-07. Liver disease has been reported as a comorbidity factor in the disease (COVID-19) caused by SARS-CoV-2 infection: PMID: 32179124, PMID:32233161. |
| GS380478 | 2550 | Cardiovascular Diseases-MeSH:D002318 | This gene set represents the genes in GS235830 queried on April 17, 2020 which was derived from the gene2mesh procedure. GS235830 was updated on 2019-01-07. Cardiovascular disease has been reported as a comorbidity factor in the disease (COVID-19) caused by SARS-CoV-2 infection: PMID: 32179124, PMID:32267833, PMID:32232218, PMID:32217556. |
| GS380470 | 1003 | Diabetes Mellitus, type 2-MeSH:D003924 | This gene set represents the genes in GS242930 queried on April 16, 2020 which was derived from the gene2mesh procedure. GS242930 was updated on 2019-01-07. Diabetes has been reported as a comorbidity factor in the disease (COVID-19) caused by SARS-CoV-2 infection: PMID: 32179124, PMID:32233161, PMID:32267833, PMID:32232218, PMID:32217650, PMID:32345579. |
| GS380479 | 1927 | Lung Diseases-MeSH:D008171 | This gene set represents the genes in GS245016 queried on April 16, 2020 which was derived from the gene2mesh procedure. GS245016 was updated on 2019-01-07. Lung (pulmonological) disease has been reported as a comorbidity factor in the disease (COVID-19) caused by SARS-CoV-2 infection: PMID:32233161. |

To create gene sets that are directly related to SARS-CoV-2 infection (COVID-19), we identified several relevant reports^5,28,29^ and captured gene sets available from these studies into the GeneWeaver environment. Details are shown in Table 2.

**Table 2:** SARS-CoV-2 (COVID-19) Gene Sets

| **Gene Set Identifier** | **Gene Count** | **Gene Set Title** | **Gene Set Description/metadata** |
| --- | --- | --- | --- |
| GS398329 | 119 | Up regulated angiogenesis and inflammation genes in lungs from patients who died from COVID-19 | This gene set describes genes that are up-regulated in lungs from patients who died from COVID-19. COVID-19 is the disease caused by SARS-CoV-2 virus. Note that this expression analysis includes only the angiogenesis-associated and inflammation-associated genes available on NanoString panels. The authors define up-regulated as those genes that show a (FDR) of $\leq 0.05$. These data are from the publication (angiogenesis) and supplementary (inflammation) materials associated with the publication. PMID:32437596 |
| GS398539 | 102 | Up regulated genes in host transcriptional response to SARS-CoV-2 in Human adenocarcinomic alveolar basal epithelial (A549) cells | This gene set describes genes that are up-regulated by the host transcriptional response to SARS-CoV-2 infection in human adenocarcinomic alveolar basal epithelial (A549) cells. COVID-19 is the disease caused by SARS-CoV-2 virus. We define up-regulated as those genes that show a (log 2 fold change) of $\geq 2$. These data are from the supplementary materials associated with the publication. PMID:32416070 |
| GS398534 | 333 | Up regulated genes in host transcriptional response to SARS-CoV-2 in Human lung adenocarcinoma epithelial (Calu3) cells | This gene set describes genes that are up-regulated by the host transcriptional response to SARS-CoV-2 infection in human lung adenocarcinoma epithelial cells derived from pleural effusion (Calu3). COVID-19 is the disease caused by SARS-CoV-2 virus. We define up-regulated as those genes that show a (log 2 fold change) of $\geq 2$. These data are from the supplementary materials associated with the publication. PMID:32416070 |
| GS398533 | 14 | Up regulated genes in host transcriptional response to SARS-CoV-2 in normal human bronchial epithelium (NHBE) cells | This gene set describes genes that are up-regulated by the host transcriptional response to SARS-CoV-2 infection in normal human bronchial epithelium cells (NHBE). COVID-19 is the disease caused by SARS-CoV-2 virus. We define up-regulated as those genes that show a (log 2 fold change) of $\geq 2$. These data are from the supplementary materials associated with the publication. PMID:32416070 |
| GS398334 | 587 | Up regulated genes in post-mortem lung samples from COVID-19-positive patients | This gene set describes genes that are up-regulated in post-mortem lung samples from COVID-19-positive patients relative to biopsied healthy lung tissue from uninfected individuals. COVID-19 is the disease caused by SARS-CoV-2 virus. We define up-regulated as those genes that show a (log 2 fold change) of $\geq 2$ . These data are from the supplementary materials associated with the publication. Note: the following HGNC id is part of this data set but was not recognized HGNC:13378.<br>PMID:32416070 |
| GS398287 | 4 | Genes that are overexpressed in severe compared to mild cases of Covid-19 | People with severe cases of Covid-19 express these proteins at significantly higher levels than people with mild cases of Covid-19. Data from Figure 2 of the paper: plasma cytokine levels in patients with COVID-19.<br>PMID:32217835 |

### Gene Set Comparison

To identify genes that were shared by all five comorbidities or four out of five comorbidities, we used the GeneWeaver ‘Combine GeneSets’ tool. To visualize the intersection of comorbidity gene sets graphically, we used the GeneWeaver ‘HiSim graph’ tool. To create a hierarchical view of interleukin pathways and the genes that are shared among comorbidities we used the HiSim graph tool at the GeneWeaver resource with homology excluded.

### Functional Analysis of Gene Sets

To evaluate the shared biology of the common genes, we interrogated the genes that were shared among comorbidities in two ways: 1) we conducted a phenotype enrichment analysis using the VLAD enrichment tool^30^ and 2) we conducted a pathway enrichment analysis using the Reactome Knowledgebase resource^31^.

#### Mammalian Phenotype Enrichment Analysis

To investigate details of the phenotypes associated with the shared genes, we took advantage of the mammalian phenotype data available from the Mouse Genome Informatics site (www.informatics.jax.org). The Mouse Genome Database (MGD) group captures phenotypic data using the Mammalian Phenotype Ontology (MP), a computable ontological structure, that can be queried and used for phenotypic enrichment analysis^32^. MGD also integrates these murine data into the context of human disease data based on orthology and gene expression^32,24^. This integrated resource allowed us to exploit the associations of mouse genes and their phenotypes for enrichment purposes using VLAD, and gives us an entry into identification of potential mouse models for future study^33^.

Mouse orthologs for the shared human genes were identified using data available from the Alliance of Genome Resources (Alliance) (www.alliancegenome.org/)^34^ using the Alliance release 3.1 stringent mouse-human orthology set. If a human gene symbol matched more than one mouse marker, that gene was not included in the analysis. This resulted in the following fourteen human genes being excluded from the analysis: *AGTR1, CCL2, CFH, CYP2D6, CYP3A4, GSTM1, GSTP1, HAMP, HLA-B, HLA-DRB1, IFNA1, MMP1, SERPINA1*, and *TIMP2*. We did not identify mouse orthologs for two human genes (*CXCL8, HLA-DQB1*). Excluding these 16 from the initial 123 genes that were shared among four of five comorbidities, left us with 107 remaining mouse orthologs which we used in the VLAD analysis. All but one of the 107 mouse genes (*H2-Ea*, a polymorphic pseudogene; i.e. a coding gene in some strains and a pseudogene in others) had annotations to MP. The VLAD phenotype analysis was run on 20 August 2020 using annotation data from 20 August 2020, ontology data from 9 July 2020 and default parameters.

#### Reactome Pathway Enrichment Analysis

For pathway enrichment, we submitted the gene lists to the Reactome ‘Analyze Gene List’ enrichment tool (https://reactome.org/PathwayBrowser/#TOOL=AT) based on Reactome version 72. The analysis was performed on 18 May 2020. Results were downloaded using the ‘Pathway Analysis Results’ and ‘Analysis Report’ functionality at Reactome.

Pathway enrichment analysis was also performed for six COVID-19-related gene sets shown in Table 2. We selected up-regulated (log2 fold change >=2) genes in host transcriptional response to SARS-CoV-2 in three cell cultures: human A549 lung alveolar cells (102 genes), Calu3 human lung adenocarcinoma epithelial cells (333 genes), normal human bronchial epithelium (NHBE) cells (14 genes); genes that are up-regulated in post-mortem lung samples from COVID-19-positive patients relative to biopsied healthy lung tissue from uninfected individuals (586 genes)^28^; immune-response and angiogenesis-related genes that are up-regulated in lungs from patients who died from COVID-19 (114 genes)^5^; genes that are overexpressed in severe compared to mild cases of COVID-19 (4 genes)^29^. We corrected for any symbols that were out of date and again used the Reactome Pathway analysis tool. The analysis was performed on 10 August 2020.

## Results

### COVID-19 comorbidities share associated genes

To test our hypothesis that comorbidities associated with COVID-19 severity have common underlying molecular bases, we chose five comorbidities that have been reported in the literature as closely associated with poorer disease outcome: Kidney Disease, Liver Disease, Diabetes, Lung Disease and Cardiovascular Disease. We searched the GeneWeaver Data repository for gene sets associated with these comorbidities and identified gene sets from MeSH and HPO that we used in our analyses (Table 1).

To identify genes that were shared among the five comorbidity gene sets, we used the ‘Combine GeneSets’ tool to create a matrix of genes and sets in which they were contained. We tabulated the number of gene sets that contained each gene and determined that eight genes were present in all five sets: *APOA1, APOE, B2M, CTLA4, F2, F5, HMOX1* and *STAT3*; 123 genes were common to at least four out of five comorbidity sets (Table 3).

**Table 3.**
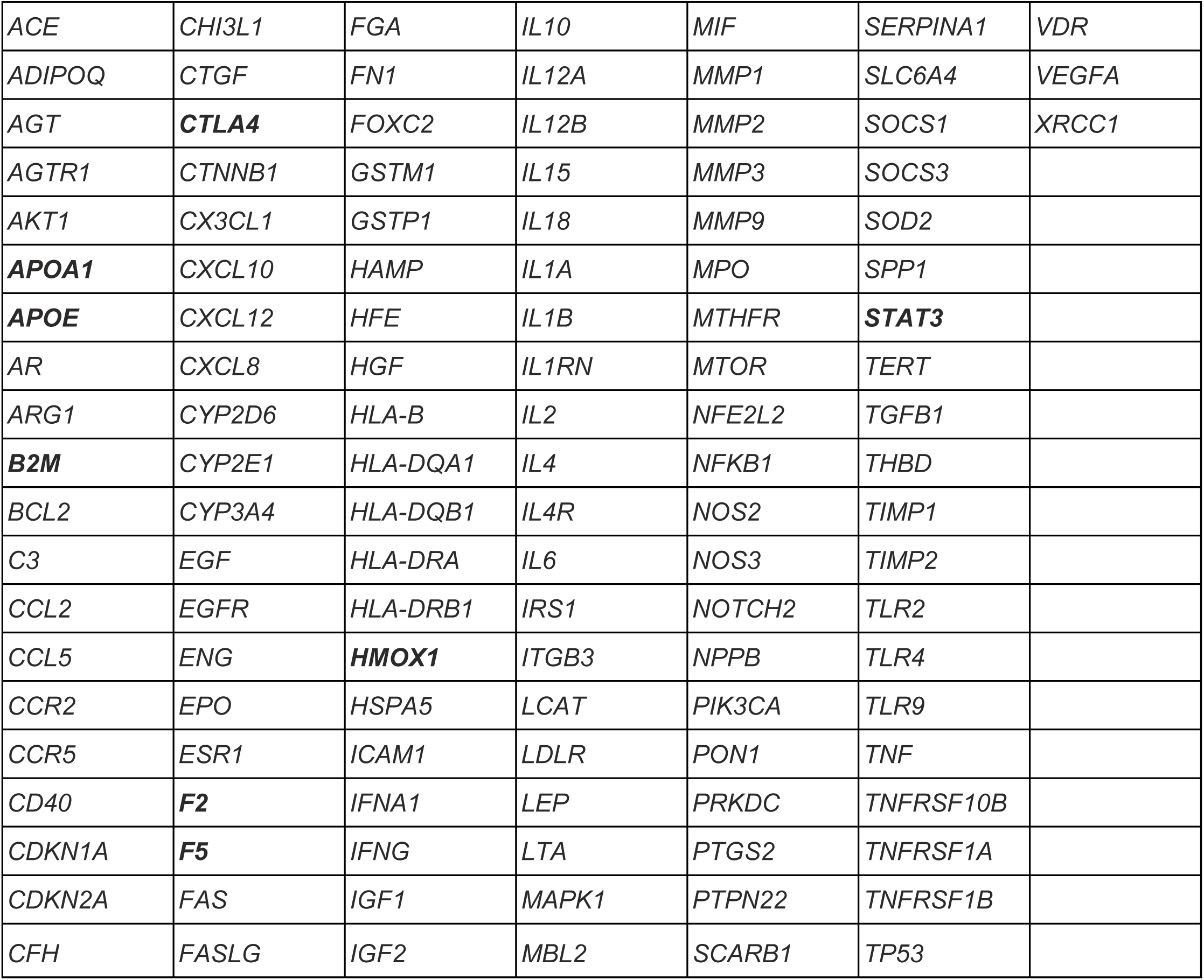
Genes shared by COVID-19 comorbidities. This table shows the genes that were annotated to four out of five comorbidities that are associated with COVID-19 severity. Genes in **bold** are annotated to all five comorbidities.

|  |  |  |  |  |  |  |
| --- | --- | --- | --- | --- | --- | --- |
| *ACE* | *CHI3L1* | *FGA* | *IL10* | *MIF* | *SERPINA1* | *VDR* |
| *ADIPOQ* | *CTGF* | *FN1* | *IL12A* | *MMP1* | *SLC6A4* | *VEGFA* |
| *AGT* | ***CTLA4*** | *FOXC2* | *IL12B* | *MMP2* | *SOCS1* | *XRCC1* |
| *AGTR1* | *CTNNB1* | *GSTM1* | *IL15* | *MMP3* | *SOCS3* |  |
| *AKT1* | *CX3CL1* | *GSTP1* | *IL18* | *MMP9* | *SOD2* |  |
| ***APOA1*** | *CXCL10* | *HAMP* | *IL1A* | *MPO* | *SPP1* |  |
| ***APOE*** | *CXCL12* | *HFE* | *IL1B* | *MTHFR* | ***STAT3*** |  |
| *AR* | *CXCL8* | *HGF* | *IL1RN* | *MTOR* | *TERT* |  |
| *ARG1* | *CYP2D6* | *HLA-B* | *IL2* | *NFE2L2* | *TGFB1* |  |
| ***B2M*** | *CYP2E1* | *HLA-DQA1* | *IL4* | *NFKB1* | *THBD* |  |
| *BCL2* | *CYP3A4* | *HLA-DQB1* | *IL4R* | *NOS2* | *TIMP1* |  |
| *C3* | *EGF* | *HLA-DRA* | *IL6* | *NOS3* | *TIMP2* |  |
| *CCL2* | *EGFR* | *HLA-DRB1* | *IRS1* | *NOTCH2* | *TLR2* |  |
| *CCL5* | *ENG* | ***HMOX1*** | *ITGB3* | *NPPB* | *TLR4* |  |
| *CCR2* | *EPO* | *HSPA5* | *LCAT* | *PIK3CA* | *TLR9* |  |
| *CCR5* | *ESR1* | *ICAM1* | *LDLR* | *PON1* | *TNF* |  |
| *CD40* | ***F2*** | *IFNA1* | *LEP* | *PRKDC* | *TNFRSF10B* |  |
| *CDKN1A* | ***F5*** | *IFNG* | *LTA* | *PTGS2* | *TNFRSF1A* |  |
| *CDKN2A* | *FAS* | *IGF1* | *MAPK1* | *PTPN22* | *TNFRSF1B* |  |
| *CFH* | *FASLG* | *IGF2* | *MBL2* | *SCARB1* | *TP53* |  |

### Genes shared among COVID-19 comorbidities are enriched for phenotypes corresponding to immune system processes and circulatory system biology

We tested the functional significance of the genes shared among the five comorbidities by performing two different kinds of enrichment analysis on our gene sets. First, we identified the mouse orthologs of the human genes and performed a phenotype enrichment using the VisuaL Annotation Display tool (VLAD) (http://proto.informatics.jax.org/prototypes/vlad/). For genes shared among all five comorbidities, VLAD enrichment analysis identified 762 significantly enriched (p=<0.05) mammalian phenotypes (supplemental table 1). The most significantly enriched terms fall into three general categories: T-cell related phenotypes, inflammation or infection related phenotypes, and cardiovascular phenotypes including blood clotting. Table 4 shows that of the eight shared genes, several were annotated to each of the significantly enriched phenotypes.

**Table 4.** Significantly enriched phenotype categories. Top significantly enriched phenotype categories identified by VLAD analysis, showing how many of the eight genes shared among all five comorbidities are annotated to each phenotype category. T Cell related phenotypes included ‘increased CD-4 positive, alpha beta T cell number’ (p=3.9×10e-8) and ‘increased T-helper Cell number’ (p=1.6×10e-06). (Complete list of enriched phenotypes available supplemental table 1)

| Phenotype Category | Gene Count | P-value range |
| --- | --- | --- |
| T Cell Related | 5 | 1.6x10e-06 to 3.9x10e-08 |
| Autoimmune Response | 5 | 1.7x10e-6 |
| Interferon gamma secretion | 4 | 3.7x10e-6 |
| Lung Inflammation | 4 | 1.7x10e-6 |
| Susceptibility to infection | 5 | 8.9x10e-7 |
| Blood Coagulation | 5 | 5.8xe10-8 |
| Cardiovascular physiology | 7 | 1.6xe10-6 |

We repeated our phenotype enrichment analysis using genes that are co-annotated to four of the five comorbidities associated with COVID-19. When we examined the shared genes among comorbidity sets, we found that 123 genes were shared among four out of five comorbidities. Phenotype enrichment analysis performed with the 107 one-to-one mouse orthologs of these human genes was consistent with our analysis of the eight genes that were conserved in all five comorbidities. The increase in gene number resulted in an increase in the number of significantly enriched mammalian phenotype terms (p=<0.05) with 3232 terms included in the enrichment analysis (supplemental table 2). VLAD analysis showed that the major areas of the ontology with the most highly significant enrichment were, as in the analysis for the eight genes shared by all the comorbidities, in inflammatory response and infection, leukocyte biology and blood vessel morphology.

Abnormal blood coagulation was no longer in the most highly significant group of phenotypes, but was significantly enriched (p=1.52×10e-11).

Similar to our results for the eight genes shared among all five comorbidities, the mouse orthologs of the 123 genes shared in four out of five comorbidities showed many genes associated with each of the most significant phenotypes (Table 5).

**Table 5.**
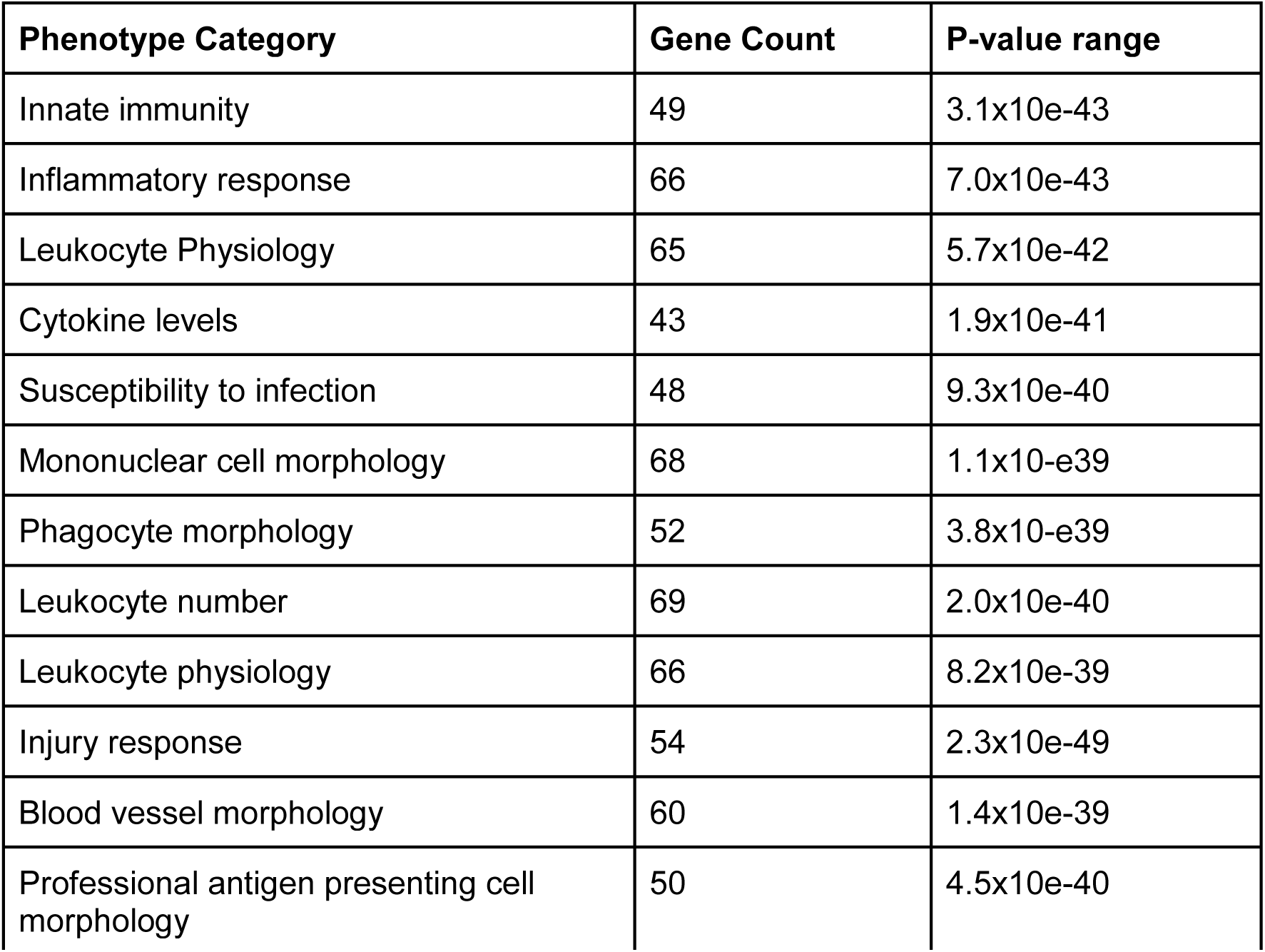
Significantly enriched phenotype categories. Top significantly enriched phenotype categories identified by VLAD analysis, showing the number of genes from the set of 107 mouse orthologs shared among four out of five comorbidities annotated to each phenotype category. (Complete list of enriched phenotypes available supplemental table 2)

### Pathway analysis enrichment includes cytokine signaling pathways, blood coagulation and plasma lipoprotein metabolism

In addition to our phenotype analysis, we were also interested in investigating whether the genes shared among comorbidities were enriched for specific biological pathways. To answer this question, we used our human gene sets and the Reactome Knowledgebase in an enrichment analysis for biological pathways.

Reactome is a manually curated resource that captures information about reactions, their relationships and the genes and chemicals that play a role in those reactions^35^.

We interrogated the Reactome Knowledgebase using the eight genes that were shared among all five comorbidities and identified 103 pathways/subpathways that were significantly enriched (FDR=<0.05, supplemental table 5). Reactome pathways are organized into a hierarchical format where grouping pathways are subcategorized into more specific pathways which in turn are eventually represented by individual reactions. Reactome captures information about not only the genes and molecules that act in a pathway but also those that are acted upon, thus casting a wide net for genes that are included in an analysis. The 25 most significantly enriched pathways grouped under several parent pathways are shown in Figure 1. Two genes, *APOA1* and *APOE*, are shared among several plasma lipoprotein assembly, remodeling and clearance pathways. Three genes, *APOA1, F2* and *F5*, are found in the hemostasis pathway, all are included in platelet activation and the latter two in blood clotting. Five of the eight genes shared among the comorbidities were involved in immune system pathways: *B2M, HMOX1, CTLA4, STAT3* and *F2*. Of these five genes, three are included in cytokine signaling: *B2M, HMOX1* and *STAT3*. Other informative pathways showed that *APOA1, APOE, F2* are in GPCR downstream signaling, and *F5, APOA1, APOE* are in vesicle-mediated transport.

**Figure 1.**
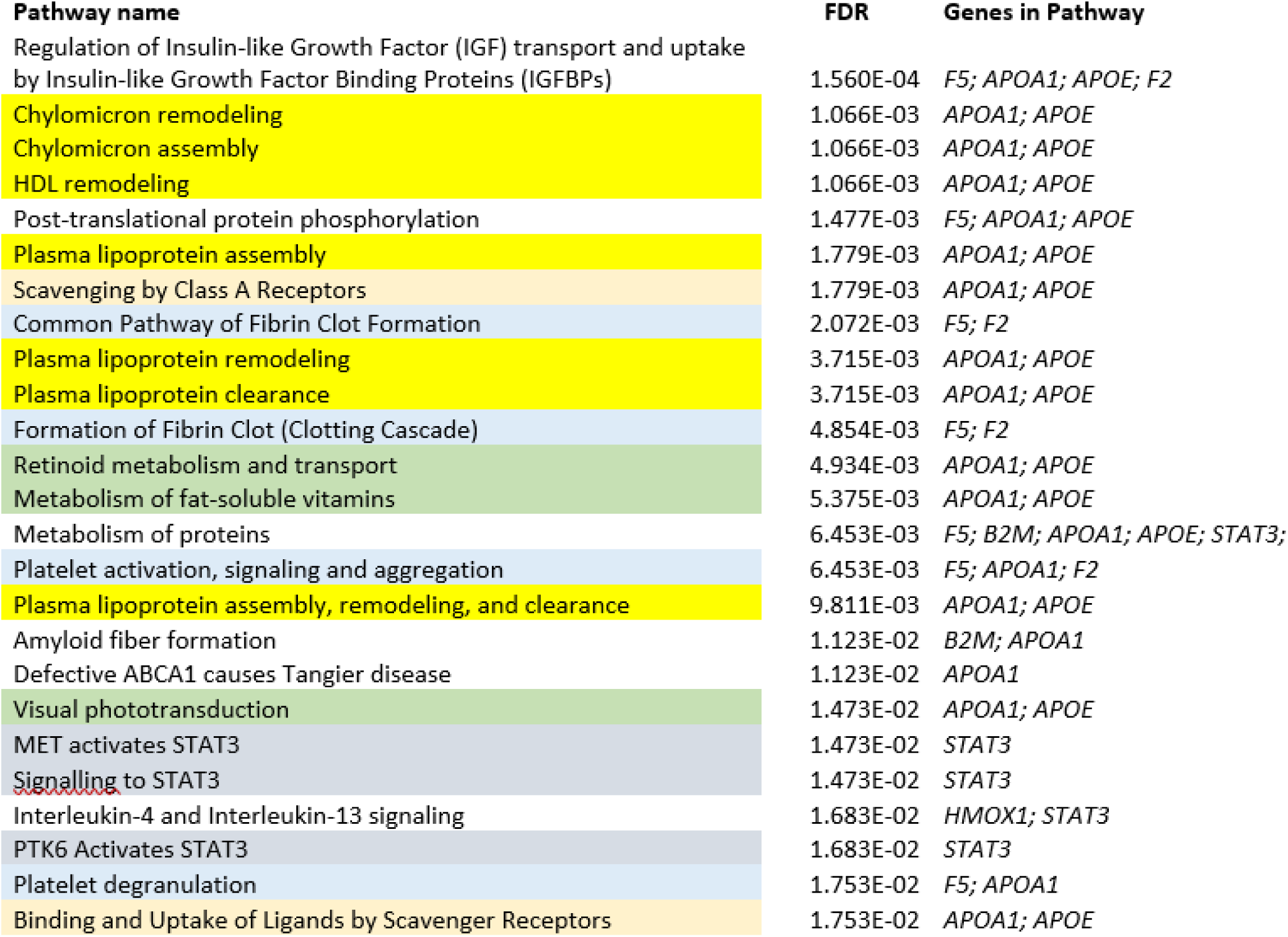
The top 25 most significantly enriched pathways involving the eight genes shared among all five comorbidities. Pathways that are similar or directly related in the Reactome knowledgebase are color coded. Yellow: lipoprotein-related processes; peach: scavenger receptor pathways; blue: blood clotting; green; retinoid-related pathways; grey; signaling through STAT3.

We repeated the pathway enrichment analysis with the 123 genes that were shared by four out of five comorbidities. We identified 172 pathways that were significantly enriched (FDR=<0.05, supplemental table 6). These results supported and confirmed the results we obtained with the eight genes that were shared among all five comorbidities. Although with lower significance, enriched pathways include ‘common pathway of fibrin clot formation’ (FDR=5.9×10e-3; four genes), ‘platelet degranulation’ (FDR=6.0×10e-6; thirteen genes) and ‘plasma lipoprotein assembly remodeling and clearance’ (FDR=0.034; five genes). Immune signaling pathways and particularly interleukin signaling pathways were frequent in our enrichment results (Figure 2). The downstream GPCR signaling pathway and the retinoid/vitamin pathways were no longer significantly enriched.

**Table 6.**
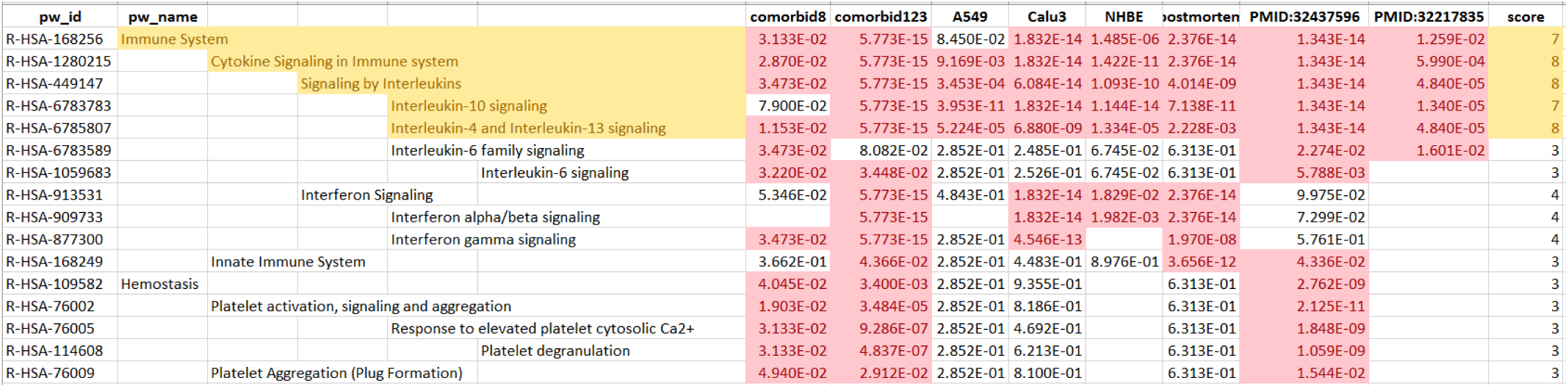
Comparison of pathway enrichment false discovery rates (FDR) for pathways with FDR <=0.05 (shown in light red) for at least one of the comorbidity sets and for at least one of the COVID-19 related gene sets: comorbid8 column displays FDR for comorbidity set for all five comorbidities; comorbid123 for comorbidity set for four of the five comorbidities; A549 for up regulated genes in host transcriptional response to SARS-CoV-2 in human A549 lung alveolar cells; Calu3 for up regulated genes in Calu3 human lung adenocarcinoma epithelial cells; NHBE for up regulated genes in normal human bronchial epithelium (NHBE) cells; postmortem for up regulated genes in post-mortem lung samples from COVID-19-positive patients; PMID:32437596 for up regulated angiogenesis and inflammation genes in lungs from patients who died from COVID-19; PMID:32217835 for genes that are overexpressed in severe compared to mild cases of Covid-19. The score column gives the number of sets with FDR meeting our criteria. Shaded in yellow are the pathways significantly enriched for both the comorbidity sets and several COVID-19 sets based on highest scores. The full table displaying 28 pathways meeting our criteria is given in supplemental material. (Complete list of enriched pathways available supplemental table 4; enriched pathway details for each gene set available supplemental tables 5-12)

**Figure 2.**
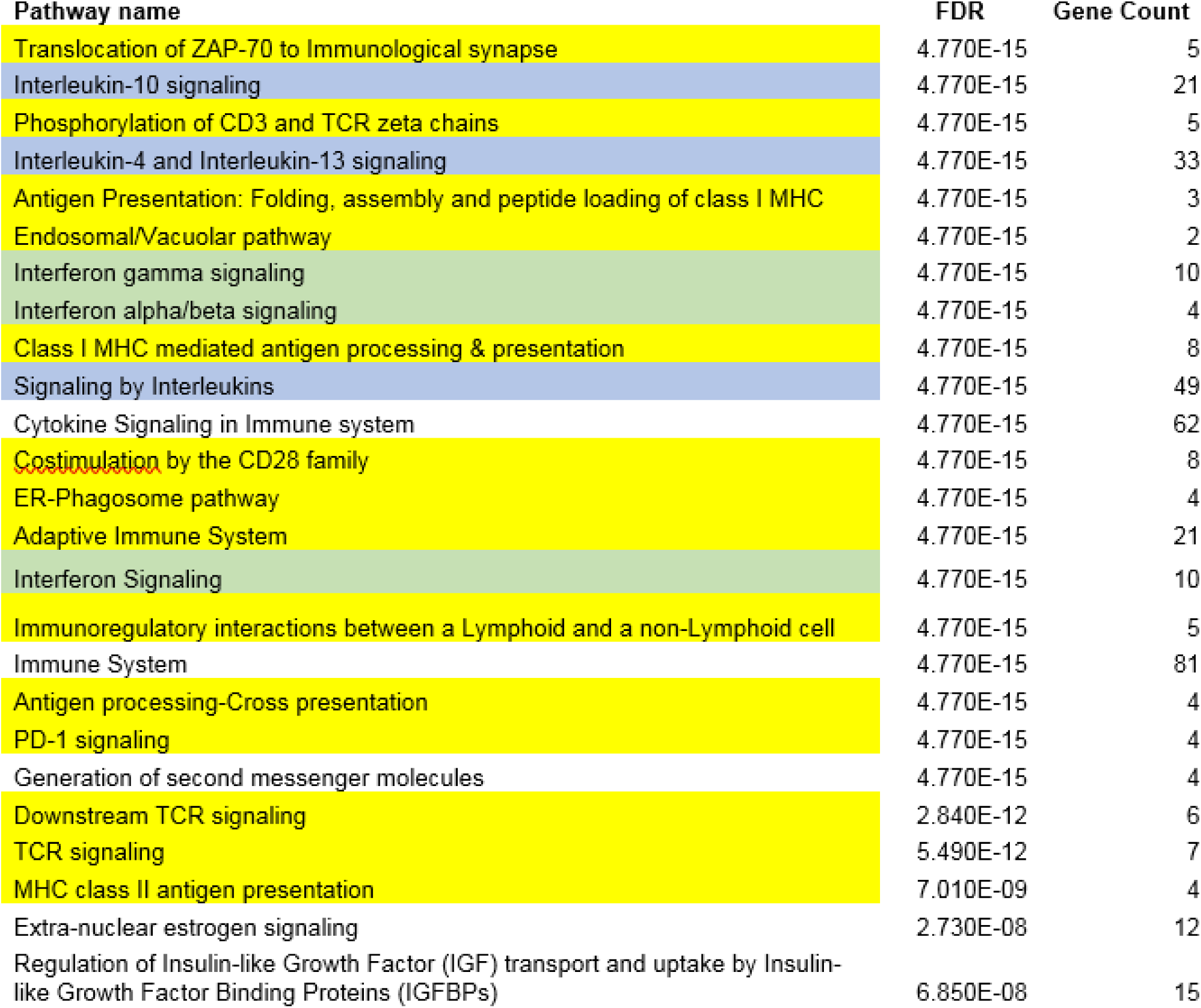
The top 25 most significantly enriched pathways involving the 123 genes shared among four out of five comorbidities. Pathways that are similar or directly related in the Reactome knowledgebase are color coded. Yellow: adaptive immune system pathways; blue: interleukin signaling pathways; green; interferon signaling pathways.

We compared our pathway enrichment results with our phenotype enrichment results for the eight genes conserved among all five comorbidities to determine if there was consistency between the results obtained from independently curated resources: MGI phenotype and Reactome. Like the pathway analysis, our phenotype enrichment analysis also revealed lipoprotein phenotypes for significant enrichment, for example ‘abnormal circulating lipoprotein level’ (p=1.81×10e-2). Phenotype analysis also revealed ‘abnormal blood coagulation’ and ‘decreased platelet aggregation’ (p=5.81×10e-8 and p=2.64×10e-2 respectively) in common with the pathway analysis. Our results from the analyses of the 123 genes conserved in four out of five comorbidities were also consistent. The pathway analysis revealed that the 25 most significant pathways were pathways related to the immune system, in particular there was concordance with the results from the eight genes, identifying pathways related to inflammatory responses, interferon and interleukin signaling. In total, the results of all of our analyses show that five comorbidities associated with severe COVID-19 share common physiological aspects including cytokine signaling, blood clotting, and plasma lipoprotein biochemistry.

### *STAT3* is shared among interleukin signaling pathways that are enriched in COVID-19 comorbidities

To further investigate whether there is a common molecular basis for the interleukin signaling pathway enrichment we examined the Reactome enrichment results using the 123 genes conserved in four out of five comorbidities for interleukin signaling pathways (supplemental table 3). We created gene sets of the shared genes that were in Reactome interleukin pathways that are significantly enriched (FDR<0.05). We used the HiSim graph tool in the GeneWeaver resource to create a graphical view of the genes that are found in the sets. The gene conserved among the largest number of sets is *STAT3* which is found in nine of the eleven significantly enriched interleukin signaling pathways. *IL12B* is shared among four signaling pathways (data not shown).

### Genes associated with SARS-CoV-2 infection response and genes shared among COVID-19 comorbidities identify common cytokine signaling pathways and hemostasis

Since we had identified pathways that were common to COVID-19-associated comorbidities, we investigated whether these pathways were also associated with the COVID-19 itself. To answer this question, we created gene sets directly associated with SARS-CoV-2 infection from published literature (Table 2). GeneWeaver Gene Set GS398287, represents four plasma cytokines that are significantly elevated in patients with severe disease versus patients with mild disease^29^. GS398329 represents 114 genes associated with angiogenesis or inflammation that were upregulated in COVID-19 postmortem samples^5^. Two gene sets, GS398539 and GS398534, of 119 and 333 genes respectively represent genes that are upregulated in two distinct lung adenocarcinoma epithelial cells infected with SARS-CoV-2; GS398533 with 14 genes represents genes that are upregulated in normal human bronchial epithelium cells infected with SARS-CoV-2; GS398334 represents 587 genes upregulated in post-mortem COVID-19 samples^28^. We ran Reactome pathway enrichment analysis on each of these sets and determined the enriched pathways that were shared with those identified in the comorbidity analyses (supplemental tables 7-12). Unsurprisingly, GS398329 and GS398287, which were preselected for genes involved in the immune response, were enriched for immune response pathways. GS398329, preselected to be associated with angiogenesis, also showed significant enrichment for the comorbidity pathways associated with platelet biology. All of the data sets showed significant enrichment for signaling mediated by interleukin-4, -10 and -13 (Table 6).

### Identification of potential mouse models to study comorbidities and COVID-19 severities

The results of our phenotype analysis using mouse orthologs of shared human genes shows that phenotypic enrichment is consistent with the pathway enrichment using the human genes and is also consistent with pathologies associated with severe COVID-19: blood coagulation, inflammation and cardiovascular pathologies^36,37,38,39,40,41^. Since mice provide an attractive genetic system for disease modeling, we investigated the phenotypes associated with each of these genes in further detail. Figure 3 shows each of the eight genes shared by all five comorbidities and the phenotype categories that were enriched in this set. Yellow highlighting indicates that mutations in the mouse gene have been annotated to a phenotype of a category that is enriched in the eight shared genes. For example homozygous mice of the genotype *Ctla4*^*tm1Shr*^/*Ctla4*^*tm1Shr*^ display multiple phenotypes that are shared with severe COVID-19: abnormal lung inflammation, abnormal cytokine secretion (interferon secretion) and autoimmune response^42^. *Hmox1*^*tm1Mlee*^*/Hmox1*^*tm1Mlee*^ mice are another example with cardiovascular, immune and liver system phenotypes (http://www.informatics.jax.org/allele/MGI:2429784).

**Figure 3.**
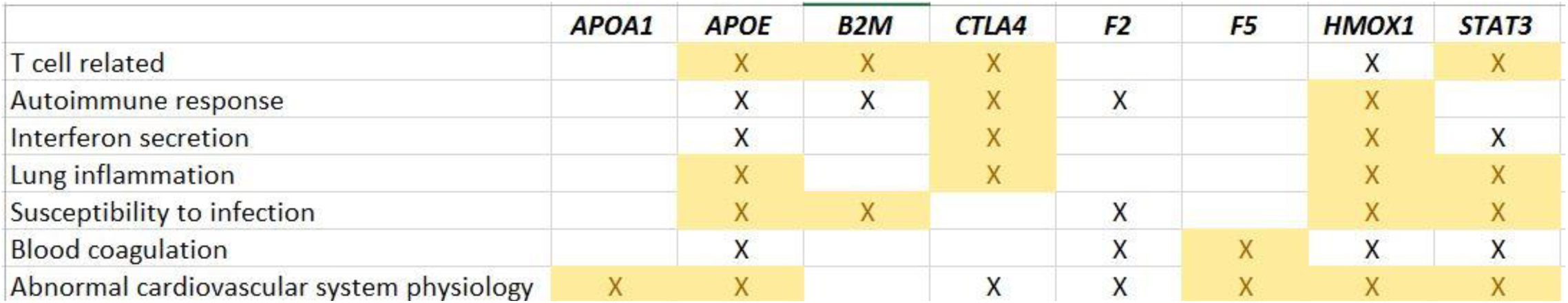
Phenotype Enrichment for the Eight Genes Shared by All Five Comorbidities. Cells shaded in yellow indicate that there is a mouse model in the MGI resource that has been studied for the specific phenotype.

## Discussion and Summary

COVID-19 is a global health concern. The disease is complex and varies in severity from asymptomatic to lethal^43^. As our understanding of the disease has progressed, a number of comorbidities associated with the disease have been identified that lead to greater severity. The goal of our work is to identify underlying genetic factors that might explain the mechanism of why certain comorbidities lead to more severe disease. To this end, we studied genetic features of five comorbidities that are associated with severe COVID-19: Cardiovascular Disease, Diabetes, Hepatitis, Lung Disease, and Kidney Disease. We identified common genes that were associated with each of the comorbidities and the pathways and phenotypes with which they are associated. We compared the results of the comorbidity analysis with genes that were directly associated with SARS-CoV-2 and showed that they shared common pathways involved in the immune response and platelet biology. Our results are encouraging in that these areas of physiology have also been correlated with severe disease. Here we discuss our results in the context of COVID-19 severity.

Our analysis of genes shared among both comorbidities and SARS-CoV-2 infection identified several interleukin signaling pathways that were enriched in both categories. Interleukin-4/-13 and interleukin-10 signaling shared enrichment among the largest number of gene sets we examined, and interleukin-6, interleukin-12 and interleukin-2 shared enrichment between at least one comorbidity set and a set of genes upregulated in patients who died from COVID-19. STAT3 positively regulates the transcription of IL-6, which controls inflammation^44^ and is a downstream signaling player in the IL-6 pathway through the IL6ST protein^45,46^. IL-12 is produced in response to infection and signals through the JAK-STAT pathway, including *STAT3*, to induce the proliferation of NK cells and T cells. These cells in turn trigger cytokine signaling including interferon gamma^47^.

One mechanism proposed for the severity of COVID-19 is the coincidence of severe acute respiratory distress triggered by a cytokine related syndrome triggered by the angiotensin signaling pathway^48^. An interesting aspect of this proposal is its action through *STAT3*, one of the genes that we also found conserved in the comorbidities we studied. Targeting the JAK-STAT pathway has been proposed as a therapeutic approach to COVID-19^2^. Our results show that *STAT3* is conserved in all five comorbidities we analyzed. This supports the hypothesis that a promising therapeutic strategy to combat severe COVID-19 compounded by preexisting comorbidities may be to target common JAK-STAT pathways.

In addition to immune signaling pathways, we also saw shared enrichment for pathways involved in platelet biology. Platelets are the cells that are responsible for blood clotting^49^. Abnormal clotting has been observed in severe COVID-19 patients and it has been suggested as a complication that leads to more severe disease^50,51^. Magro *et al*. reported that the abnormal clotting observed in severe COVID-19 patients correlated with activation of the complement pathway^52^. Our results show that *HMOX1, APOA1, APOE* and two members of the coagulation cascade, *F2* and *F5*, are shared among all five comorbidities we examined. In mice, *Hmox1* deficiency leads to coagulation defects and results in arterial damage due to oxidative stress^53^. APOA1 is released during platelet degranulation as part of the platelet secretory granule^54^ (Reactome:R-HSA-482770). APOA1 levels have also been shown to be significantly decreased in severe COVID-19 patients^55^. *APOE* is a gene that encodes a lipid binding protein involved in cholesterol metabolism^56^. Preliminary evidence suggests that the e4 allele of *APOE* may lead to a higher risk of deep vein thrombosis and the same allele also predicts severe COVID-19^57,58^. *F2* and *F4* are both involved in the formation of a fibrin clot (Reactome:R-HSA-140877). The complement pathway and coagulation cascade have been shown to interact, tying together inflammation and hemostasis^59^. Additionally in a proteomic study of proteins differentially expressed in the serum of 28 severe COVID-19 patients compared with non-severe COVID-19 patients, Shen *et al*. report that 50 of 93 differentially regulated proteins fall into three categories one of which is platelet degranulation^55^. These results suggest that one of the factors contributing to severe disease in patients with any of the five comorbidities may be due to an underlying genetic mechanism that acts through the hemostatic pathway.

Our results show that genes that are shared among five comorbidities associated with severe COVID-19 identify pathways that are consistent with the pathologies associated with the disease. In our analysis we excluded mouse orthologs that did not correlate 1:1 with human genes to avoid potential skewing of the enrichment analysis by having multiple paralogs over-represented. Despite this, and the exclusion of several potentially important immune system genes such as some histocompatibility genes, our results show that analysis using mouse orthologs of the shared genes also identifies phenotypes that are consistent with disease pathology. The laboratory mouse provides a tractable system to study the effects of genetic foundations of the comorbidities and severe disease. As mentioned above mice carrying *Ctla4*^*tm1Shr*^/*Ctla4*^*tm1Shr*^ and *Hmox1*^*tm1Mlee*^*/Hmox1*^*tm1Mlee*^ homozygous mutations display multiple phenotypes consistent with severe COVID-19 pathology. Mice engineered for mutations in these genes crossed with mice engineered to carry the human *ACE-2* SARS-CoV-2 viral receptor, ICR-Tg(Ace2-ACE2)1Cqin/J, would be a starting point to explore the underlying genetic variants related to comorbidities interact with viral infection^60^. Humanized mice, like the *Apoe*^*tm3*(*APOE*\**4*)*Mae*^ strain, which carries the human E3 variant mentioned above as being implicated in both severe disease and an underlying thrombosis pathology, could be used in conjunction with ICR-Tg(Ace2-ACE2)1Cqin/J to study the effects of the human variant on viral infection. A comprehensive resource for using the mouse as a model system for COVID-19 research is maintained by the Mouse Genome Informatics Group [http://www.informatics.jax.org/mgihome/other/coronavirus.shtml].

In this study we have used a bioinformatics approach to interrogate genes associated with five COVID-19 comorbidities that correlate with severe disease. Using genes that have been annotated to these comorbidities in the MeSH or HPO resource we have shown that genes are shared among the comorbidities and that shared genes are enriched for pathways that could be the genetic basis for the pathologies observed with severe COVID-19, specifically our results suggest that the interrelated pathways of hemostasis and inflammation may be key players in understanding the severity of comorbidities with COVID-19^61,62^. Our studies provide a gateway to understand how host genetics interacts with and influences the consequences of viral infection. Our knowledge about COVID-19 continues to grow at a rapid rate and future work will entail the examination of additional comorbidities, more specific comorbidities, a wider survey of genes beyond our initial seed set from MeSH and HPO. As we learn more about correlations between individual comorbidities and disease pathologies, we may be able to identify specific pathway/comorbidity combinations that can be used to inform us about treatment decisions. Our work also provides an entry point into an experimental system using the laboratory mouse to manipulate host genetics and to study its subsequent effect on the pathology of viral infection.

## Data availability

All gene sets generated during and analyzed during the current study are based on data published in peer-reviewed papers, are available in the public GeneWeaver repository [www.geneweaver.org] and are accessible using the gene set identifiers given in the text (e.g. GS398287). Results data generated during this study are included in this published article and its supplementary files.

## Acknowledgements

This work was funded by NIH grants to the Mouse Genome Database (NHGRI U41 HG000330), the Jackson Laboratory Center for Precision Genetics (OD U54 OD020351) and GeneWeaver (NIAID RO1 AA18776). The authors would like to thank Dr. Peter D’Eustachio and Dr. Laurens Wilming for their critical reading of the manuscript.

## Author contributions

DPH and JAB conceived the study and were in charge of overall direction and planning. DPH and MED carried out the implementation and performed the computations. GM, MSM, and EJC contributed to the interpretation of the results. DPH took the lead in writing the manuscript with support from MED. All authors discussed the results and contributed to the final manuscript.

## Author affiliations

Mary E. Dolan, David P. Hill, Gaurab Mukherjee, Monica S. McAndrews, Elissa J. Chesler, Judith A Blake The Jackson Laboratory, Bar Harbor, ME 04609, USA

## Materials & Correspondence

Correspondence and material requests should be addressed to Mary E. Dolan.

